# mTOR inhibition enhances delivery and activity of antisense oligonucleotides in uveal melanoma cells

**DOI:** 10.1101/2021.09.29.462324

**Authors:** Shanna Dewaele, Louis Delhaye, Boel De Paepe, Bram Bogaert, Ramiro Martinez, Jasper Anckaert, Nurten Yigit, Justine Nuytens, Rudy Van Coster, Sven Eyckerman, Koen Raemdonck, Pieter Mestdagh

**Affiliations:** OncoRNALab, Center for Medical Genetics (CMGG), Ghent University, Ghent, 9000, Belgium; Cancer Research Institute Ghent (CRIG), Ghent, 9000, Belgium; Department of Pediatrics, Division of Pediatric Neurology and Metabolism, Ghent University Hospital, Ghent, 9000, Belgium; Center for Medical Biotechnology, VIB-Ghent University, Ghent, 9052, Belgium; Department of Biomolecular Medicine, Ghent University, Ghent, 9000, Belgium; Laboratory for General Biochemistry and Physical Pharmacy, Ghent University, Ghent, 9000, Belgium

## Abstract

Uveal melanoma (UM) is the most common primary intraocular malignancy in adults. Due to a lack of effective treatments, patients with metastatic disease have a median survival time of 6-12 months. We recently demonstrated that the *SAMMSON* long non-coding RNA (lncRNA) is essential for uveal melanoma cell survival and that antisense oligonucleotide (ASO)-mediated silencing of *SAMMSON* impaired cell viability and tumor growth *in vitro* and *in vivo*. By screening a library of 2911 clinical stage compounds, we identified the mTOR inhibitor GDC-0349 to synergize with *SAMMSON* inhibition in UM. Mechanistic studies revealed that mTOR inhibition enhanced uptake and reduced lysosomal accumulation of lipid complexed *SAMMSON* ASOs, improving *SAMMSON* knockdown and further decreasing UM cell viability. We found mTOR inhibition to also enhance target knockdown in other cancer cell lines as well as normal cells when combined with lipid nanoparticle complexed or encapsulated ASOs or small interfering RNAs (siRNAs). Our results are relevant to nucleic acid treatment in general and highlight the potential of mTOR inhibition to enhance ASO and siRNA mediated target knockdown.

## Introduction

Uveal melanoma is a genetically and biologically distinct type of melanoma harboring chromosomal aberrations which includes loss of 1p, monosomy of chromosome 3, loss of 6q and 8p and gain of 6p and 8q as well as mutations in *GNAQ* or *GNA11* (90% of UM tumors)(1). While rare, it is the most common intra-ocular malignancy in adults with an incidence of 5 - 7 cases per million annually(2, 3). UM tumors develop in the iris of the eye (anterior UM) or in the choroid and/or ciliary body (posterior UM). Approximately 90 - 95% of UM cases are posterior UM and are characterized as highly aggressive and frequently metastasizing(4, 5). Despite successful local treatments, UM tumors metastasize to the liver, lungs or bones in 50% of the UM patients(6). Currently, no effective treatments are available for metastatic uveal melanoma, resulting in a median survival time of 6-12 months(5, 7).

Long non-coding RNAs (lncRNAs) are a well characterized class of non-coding RNAs (ncRNAs), defined as >200 nucleotide long transcripts without protein-coding potential(8). The lncRNA Survival Associated Mitochondrial Melanoma Specific Oncogenic Non-coding RNA (*SAMMSON*) was originally discovered on chromosome 3p13 as a lineage survival oncogene in skin melanoma(9) and further validated as a candidate therapeutic target in UM and conjunctival melanoma (CM) cells(10). Antisense oligonucleotide (ASO)-mediated *SAMMSON* knockdown in UM cells impairs protein synthesis and mitochondrial function, resulting in a potent anti-tumor response *in vitro* and *in vivo*(10).

ASOs and small interfering RNAs (siRNAs) are promising therapeutic tools to target RNA and modulate protein expression of disease-associated genes. Despite their great potential, the delivery to specific tissues and cellular uptake poses a major challenge and limitation(11, 12). Decades of research have improved the stability and biological activity of ASOs and siRNAs by adding chemical modification of the ribose group, including 2’-O-methyl (2’-OMe), 2’-O-(2-methoxyethyl) (2’-MOE), locked nucleic acid (LNA) and constraint ethyl (cET) in combination with a phosphorothioate (PS) backbone(13–15). Although several studies described the capability of cell lines to freely take up ASOs, many cell lines including uveal melanoma cell lines lack this capacity(16–18). Improved ASO and siRNA delivery to specific organs such as the liver has been achieved by conjugating a tris N-galactosamine (GalNAc) domain which binds to trimeric asialoglycoprotein receptors (ASGPRs) that are highly abundant on hepatocytes(19). A more general approach is the use of viral and non-viral vectors as ASO and siRNA delivery systems. Compared to viral vectors, non-viral vectors including lipid nanoparticles (LNPs) and polymer nanomaterials are considered safer, easier in production and are capable of carrying larger payloads(20). Since the approval of siRNA drug ONPATTRO^®^ (patisiran), LNP-based delivery is amongst the most widely used delivery systems(21). Although delivery systems enhance circulation lifetimes and improve extravasation to disease sites, site-specific bioavailability is rather limited by intracellular biological barriers including lysosomal entrapment(22, 23). Several studies have been conducted to identify compounds that can overcome these barriers by enhancing the cellular uptake and/or endosomal escape (e.g. siRNA loaded nanogels and cholesterol-conjugated siRNAs in combination with cationic amphiphilic drugs (CADs)(24–27) and siRNAs in combination with Guanabenz acetate(28)). Other studies identified specific proteins involved in the trafficking and maturation of endolysosomal compartments, and by modulating their expression, escape from these compartments can be facilitated(29).

Here we combined *SAMMSON* inhibition with a library of clinical stage compounds to identify synergistic combinations impairing uveal melanoma cell proliferation. We identified mammalian target of rapamycin (mTOR) inhibition to synergize with lipid assisted ASO-mediated *SAMMSON* inhibition through an enhancement of ASO uptake and reduced lysosomal accumulation and consequently *SAMMSON* knockdown. Lipid nanoparticle-based delivery of ASOs and siRNAs could also be enhanced in other cell types when combined with mTOR inhibition.

## Material and methods

### Cell culture and generation of stable cell lines

For all human cell lines, an ethical approval was obtained from the Ghent University commission for medical ethics. The human uveal melanoma cell lines 92.1(30) and OMM1(31) were obtained from the Leiden University Medical Center, The Netherlands. Lung adenocarcinoma A-549 cells were obtained from the Laboratory of Pharmaceutical microbiology, Belgium. Bronchial epithelial cells BEAS-2B and human embryonic kidney cells HEK293T were obtained from ATCC (Manassas, Virginia, USA). Neuroblastoma cell line SK-N-BE(2)-C was obtained from Northern Institute for Cancer Research, United Kingdom. The UM cell lines were grown in Dulbecco’s modified Eagle’s medium (DMEM, Gibco)/F12 – Roswell Park Memorial Institute (RPMI, Gibco) 1640 (1:1) medium. A-549 cells were grown in Ham’s F-12K (Kaighn’s) medium (Gibco) and HEK293T in DMEM. BEAS-2B and SK-N-BE(2)-C cells were grown in RPMI 1640. All media were supplemented with 10% fetal bovine serum (FBS), 2 mM L-glutamine (Gibco) and 100 IU/ml penicillin/streptomycin (Gibco) and the cell lines were incubated in a humidified atmosphere containing 5% CO_2_ at 37°C. For conducting experiments, cells were grown in L-glutamine and penicillin/streptomycin free media. Short tandem repeat (STR) genotyping was used to validate cell line authenticity and absence of mycoplasma was verified on a monthly basis for all cell lines in culture.

For the generation of UM cell lines with stable GFP expression, pLV-EFS-NLS-msfGFP-mAID_mPGK-BlastR was build using an in-house modified Golden GATEway(32) cloning system that was adapted for mammalian expression vectors. The construct pLV-EFS-NLS-msfGFP-mAID_PGK-BlastR was transformed and grown in DH5a competent cells, and plasmid DNA from a 250 ml culture was isolated with the Nucleobond Xtra Midi prep (Macherey Nagel, Allentown, Pennsylvania, USA, catalog #MN 740410.50). Subsequently, the plasmid was transfected and packaged in HEK293T using a second-generation lentiviral system consisting of the pCMVR7.84 packaging vector (Addgene, Watertown, Massachusetts, USA, catalog #22036) and the pMD2.G VSV-G-expressing envelope vector (Addgene, catalog #12259) to produce pseudotyped lentivirus. Transfection was performed with TransIT-LT1 (Mirus, Madison, Wisconsin, USA) according to manufacturer’s instructions. The virus was harvested after 48 h, and the supernatant was filtered, titrated, and stored at −80 °C until further use. UM cells were transduced with virus supplemented with 8 µg/ml polybrene (Sigma Aldrich, Saint Louis, Missouri, USA) and 24 h after transduction, cells were refreshed with complete medium. Transduced cells were selected after 48 h using 6 µg/ml Blasticidine S (Sigma Aldrich) for 2 weeks.

### Compounds, antisense oligonucleotides (ASOs) and small interfering RNAs (siRNAs)

The 2911 compounds used in the screening were obtained from the Centre for Drug Design and Discovery (CD3, KU Leuven, Belgium). For validation experiments, the compounds were purchased separately. Amsacrine was purchased from MedChemExpress (Monmouth Junction, New York, USA). Pimozide was purchased from Sigma Aldrich. CI-994, Clioquinol, ebastine, entinostat, fenofibric acid, fluphenazine dihydrochloride, GDC-0349, INH-6, LDN-193189 2HCl, mifepristone, oltipraz (4-methyl-5-pyrazin-2-yldithiole-3-thione), oxolinic acid, perphenazine, propylthiouracil, ranitidine hydrochloride, riboflavin, temsirolimus, torkinib and rapamycin were purchased from Selleckchem (Houston, Texas, USA).

The LNA GapmeR oligonucleotides specifically targeting *SAMMSON* (*SAMMSON* ASO), *MALAT1* and *GAPDH* and non-targeting control (NTC) GapmeR were purchased from Qiagen (Hilden, Germany).

Sequence LNA *SAMMSON* ASO*: GTGTGAACTTGGCT (catalog number 339517)

Sequence LNA NTC ASO: TCATACTATATGACAG (catalog number 339517)

Sequence LNA MALAT1 ASO: CGTTAACTAGGCTTTA (catalog number 339515)

Sequence LNA GAPDH ASO: AGATTCAGTGTGGTGG (catalog number 339515)

Sequences eGFP siRNA**: Sense strand (5’>3’): CAAGCUGACCCUGAAGUUCtt, Antisense strand (5’>3’): GAACUUCAGGGUCAGCUUGtt

Sequences NTC siRNA**: Sense strand (5’>3’): UGCGCUACGAUCGACGAUGtt, Antisense strand (5’>3’): CAUCGUCGAUCGUAGCGCAtt

*: for confocal imaging, *SAMMSON* ASO is labeled with FAM at the 5’ end

**: capital and lower case letters represent ribonucleotides and 2’-deoxyribonucleotides, respectively

### Synthesis and physicochemical characterization of ASO-loaded MC3 Lipid Nanoparticles

MC3 ASO and siRNA LNPs were synthesized as described in previous work by Van de Vyver et.al.(26) by injecting one volume of lipid mixture of DLin-MC3-DMA ((6Z,9Z,28Z,31Z)-heptatriaconta-6,9,28,31-tetraen-19-yl4- (dimethylamino)butanoate, abbreviated as MC3), DSPC (1,2-distearoyl-sn-glycero-3-phosphocholine), cholesterol and DMG-PEG2000 (1,2-dimyristoyl-rac-glycero-3-methoxypolyethylene glycol-2000) (50:10:38.5:1.5 mol ratio) in ethanol and three volumes of ASO or siRNA (optimal molar N/P charge ratio of 5) in acetate buffer (pH 5, 10 mM) in the microfluidic NanoAssemblr® Benchtop mixing device (Precision Nanosystems, Vancouver BC, Canada) at a total flow rate of 12 mL/min (3 mL/min for ethanol and 9 mL/min for aqueous buffer, flow rate ratio of 3:1 (aqueous to ethanol)). The resultant mixture (5.8 mg/mL total lipid concentration) was dialyzed (Pur-A-Lyzer™ Maxi 12000 Dialysis Kit) overnight against phosphate buffered saline (PBS) to remove residual ethanol and to raise the pH to 7.4. DLin-MC3-DMA was purchased from MedChemExpress. All other lipids were purchased from Avanti Polar Lipids, Inc. (Alabaster, Alabama, USA). Samples were stored at 4 °C until use. Hydrodynamic diameter, zeta-potential and polydispersity index (PDI) of the MC3 ASO and siRNA LNPs (after dialysis) were determined in HEPES buffer (pH 7.4, 20 mM) via Dynamic Light Scattering (DLS) (Zetasizer Nano, Malvern Instruments Ltd., Worcestershire, United Kingdom). ASO/siRNA complexation and encapsulation efficiency were respectively determined by a Quant-iT™ RiboGreen® RNA assay (ThermoFisher Scientific).

### ASO-compound screening

Before conducting the initial compound screening, the therapeutic response window was delineated by testing various concentrations of the *SAMMSON* ASO in combination with various concentrations of the compound cycloheximide of which is known to affect cell viability. Consequently, a fixed concentration of both *SAMMSON* ASO (35 nM) and compounds (10 µM) was used for the initial screening. For the ASO-compound screening, UM cell line 92.1 was seeded in opaque 384 well plates (Greiner Bio-one, Vilvoorde, Belgium, cat#788010) at a density of 1250 cells/well. After overnight incubation, the cells were transfected using lipofectamine 2000 (ThermoFisher Scientific, Waltham, Massachusetts, USA) with 35 nM *SAMMSON* ASO or NTC ASO using a Multidrop Combi reagent Dispenser (Thermo Fisher Scientific). From each compound plate, compounds were added using the Janus Mini MDT workstation (PerkinElmer, Waltham, Massachusetts, USA) to a *SAMMSON* ASO transfected plate and an NTC transfected plate, to a final compound concentration of 10 µM and final DMSO concentration of 0.1%. Cell viability was examined 48 h after treatment using a CellTiter-Glo assay (Promega, Madison, Wisconsin, USA). Before initiating the assay, the culture plates and reconstituted assay buffer were placed at room temperature for 15 minutes. Next, 20 µl of CellTiter-Glo reagent was added per well and plates were incubated for 15 minutes at room temperature to induce cell lysis. The luminescence signal was measured with a Envision plate reader (PerkinElmer). All combinations were performed on multiple plates and to compare results, intraplate normalization via two-way median polish and interplate normalization via B-score calculation was conducted. B-scores represent the effect of a compound on the viability compared to all samples in the same condition(33). The confirmation experiment was performed as described above, but included 4 replicates per condition.

### ASO-compound validation

For the validation experiments, UM cell lines 92.1 and OMM1 were seeded in 96 well plates (Corning, Glendale, Arizona, USA, costar 3596) at a density of 5000 cells/well and were allowed to settle overnight. Subsequently, the cells were transfected using lipofectamine 2000 with 50 or 100 nM LNA *SAMMSON* ASO or LNA NTC and treated with various concentrations of the compounds and with a final DMSO concentration of maximum 0.1%. The compounds were added to the wells using the HP D300e Digital dispenser (Tecan, Männedorf, Switzerland). Control cells were transfected with LNA NTC ASO and a maximum of 0.1% DMSO was added.

Cell viability was examined using a CellTiter-Glo assay (Promega). Before initiating the assay, the culture plates and reconstituted assay buffer were placed at room temperature for 30 minutes. Next, the culture medium was replaced by 200 µl fresh culture medium - assay buffer (1:1) mix. To induce complete cell lysis, the plates were shaken for 10 min. 100 µl from each well was subsequently transferred to an opaque 96-well plate (Nunc), which was measured with a GloMax 96 Microplate Luminometer (Promega). The CellTiter-Glo assay was performed on various predefined timepoints.

For real-time analysis, the IncuCyte Zoom system and IncuCyte S3 system (Essen BioScience, Newark, England) were used. Cells were seeded and treated as described above. After treatment, the culture plate was incubated in an IncuCyte Zoom system or IncuCyte S3 system at 37°C in a humidified 5% CO2 incubator. Phase contrast whole well images were captured every 3 h. The IncuCyte ZOOM (version 2016B) and IncuCyte S3 software (version 2019B) (Essen BioScience) were utilized in real-time to measure % confluence, as a proxy for proliferation. The % confluence values were corrected for seeding variability resulting in normalized % confluence values.

### Monitoring of synergistic ASO-compound combinations

Bliss independence (BI) scores were calculated to determine ASO-compound interactions. The predictive performance (BI score) is then compared to the observed drug interaction data measured by the Excess over Bliss. Excess over Bliss scores > 0 indicate drug synergism, Excess over Bliss scores = 0 indicate additive effects and Excess over Bliss scores < 0 indicate drug antagonism.

### SUnSET

Cells were seeded in T75 culture flasks (Greiner Bio-One, Cellstar) at a density of 1.17 × 10^6^ cells/flask 24h prior to transfection. The cells were transfected with 50 nM *SAMMSON* ASO or NTC ASO using lipofectamine 2000 and subsequently treated with 0.625 µM of GDC-0349 or 0.006% DMSO. After 24h, cells were washed in 1x phosphate buffer saline (PBS) and subsequently incubated with puromycin containing media (InvivoGen, San Diego, California, USA, 10 µg/ml) for 10 min. Puromycin incorporation is a proxy for the mRNA translation rate *in vitro* and was measured by western blotting using an anti-puromycin antibody (MABE343, clone 12D10, Merck Millipore, Burlington, Massachusetts, USA, 1:10 000). The antibody was diluted in Milk/TBST (5% non-fat dry milk in TBS with 0.1% Tween20). A ponceau S staining (Sigma Aldrich, Saint Louis, Missouri, USA) was performed to verify equal loading.

### Western blot analysis

Cells were lysed in RIPA lysis buffer (5 mg/ml sodium deoxycholate, 150 mM NaCl, 50 mM Tris-HCl pH 7.5, 0,1% SDS solution, 1% NP-40) supplemented with protease and/or phosphatase inhibitors. Protein concentrations were determined with the BCA protein assay (Bio-Rad, Hercules, California, USA). In total, 35 μg of protein lysate was loaded onto an SDS-PAGE gel (10% Pre-cast, Bio-Rad), ran for 1 h at 100 V and subsequently blotted onto a nitrocellulose membrane. HRP-labeled anti-rabbit (7074 S, Cell Signaling, Danvers, Massachusetts, USA, 1:10000 dilution) and anti-mouse (7076P2, Cell Signaling, 1:10000 dilution) antibodies were used as secondary antibodies. The antibodies were diluted in Milk/TBST (5% non-fat dry milk in TBS with 0,1% Tween20) and antibody binding was evaluated using the SuperSignal West Dura Extended Duration Substrate (ThermoFisher Scientific) or SuperSignal West Femto Maximum Sensitivity Substrate (ThermoFisher Scientific). Imaging was done using the Amersham Imager 680 (GE Healthcare, Chicago, Illinois, USA). Image J (version 1.52q) was used for the quantification of the blots. Uncropped scans of the blots can be found in Supplemental Fig 3.

### Seahorse XF Cell Mito Stress Test

A seahorse XF Cell Mito Stress Test was performed to measure the oxygen consumption rate (OCR). Cells were seeded in T75 culture flasks (Greiner Bio-One, Cellstar) at a density of 1 170 000 cells/flask 24 h prior to transfection. The cells were transfected with 50 nM *SAMMSON* ASO or NTC ASO using lipofectamine 2000 and subsequently treated with 0.625 µM of GDC-0349 or 0.006% DMSO. Four hours later, 25 000 cells were transferred to Seahorse XFp Cell Culture Miniplates (Agilent Technologies, Santa Carla, California, USA) and were allowed to settle overnight. Subsequently, oxygen consumption rates were measured in triplicates for each condition using the Seahorse XFp device (Agilent Technologies) according to the standard mito stress test procedures in seahorse assay medium supplemented with 14.3 mM glucose, 1 mM pyruvate and 2 mM glutamine (Sigma Aldrich), and cells were sequentially challenged with 1 µM oligomycin, 0.5 µM (92.1) or 1 µM (OMM1) carbonyl cyanide 4-(trifluoromethoxy)-phenylhydrazone (FCCP) and 0.5 µM of a rotenone antimycin A mix (Agilent Technologies). Following the assay, protein concentrations were calculated based upon absorbance reading at 280 nm (Biodrop, Isogen Life Science, Utrecht, The Netherlands) for normalization of the results. Spare respiratory capacity was calculated as the difference between maximal and the basal oxygen consumption rate. Relative oxygen consumption rates are relative compared to NTC ASO values.

### Confocal microscopy

Cells were seeded on coverslips in 24 well plates at a density of 30 000 cells/well and were allowed to settle overnight. Subsequently, the cells were transfected using lipofectamine 2000 (Life Technologies) with 50 nM *SAMMSON* ASO 3’ FAM or NTC ASO and treated with 0.625 µM of GDC-0349 or 0.006% DMSO. After 24h, cells were incubated with LysoTracker Red DND-99 (ThermoFisher Scientific, L7528, 1 µM) for 2 h, followed by washing one time with 1x PBS. Cells were fixed with 4% paraformaldehyde for 25 min at room temperature and washed 3 times with PBS. After incubation with Hoechst 33342 (ThermoFisher Scientific, H3570, 1:2000 dilution in PBS) for 10 min at room temperature, cells were washed 3 times with PBS and permeabilized at room temperature for 15 min with 0.1% Triton X-100 (Sigma Aldrich). Next, cells were washed 3 times with PBS and stained with HCS CellMask Deep Red (ThermoFisher Scientific, H32721, 1:5000 dilution in PBS) for 30 min at room temperature. After washing the cells 3 times with PBS, coverslips were transferred to a microscopic glass slide. A drop of 1% propylgallate in glycerol was used as mounting medium and coverslip sealant before acquiring images using a confocal microscope (Zeiss LSM880 with AiryScan, Jena, Germany). Five random fields per condition were selected with a total of at least 30 cells per condition. Z-stacks were generated from images taken at 0.19 µm depth per section. Volumes of the entire cell, lysosomes, nucleus and ASOs within the cell compartments were calculated using Volocity (PerkinElmer, version 6.3).

### Reverse transcription quantitative polymerase chain reaction (RT-qPCR)

Total RNA was extracted using the miRNeasy kit (Qiagen) according to the manufacturer’s instructions, including on-column DNase treatment. The Nanodrop (ThermoFisher Scientific) was used to determine RNA concentrations and cDNA synthesis was performed using the iScript Advanced cDNA synthesis kit (Bio-Rad) using a mix containing 200 ng of RNA, 4 µl of 5x iScript advanced reaction buffer and 1 µl of iScript advanced reverse transcriptase. The qPCR reactions contain 2 µl of 1:4 diluted cDNA (2.5 ng/µl), 2.5 µl SsoAdvanced Universal SYBR Green Supermix (Bio-Rad), 0.25 µl forward (5 µM, IDT, Coralville, Iowa, USA) and 0.25 µl reverse primer (5 µM, IDT) and was analyzed on a LC480 instrument (Roche, Basel, Switzerland).

When cells were seeded and treated in 96-well plates, RNA was obtained using the SingleShot Cell Lysis Kit (Bio-Rad) according to the manufacturer’s instructions and cDNA synthesis was performed using the iScript Advanced cDNA synthesis kit (Bio-Rad) using a mix containing 8 µl sample lysate, 7 µl nuclease free water, 4 µl of 5x iScript advanced reaction buffer and 1 µl of iScript advanced reverse transcriptase. Subsequently, the cDNA is diluted 4 times prior to the qPCR reaction (see higher). Expression levels were normalized using expression data of at least 2 stable reference genes out of 4 tested candidate reference genes (SDHA, HPRT1, UBC and TBP). Multi-gene normalization and relative quantification was performed using the qbase+ software (v3.2, www.qbaseplus.com).

The primer sequences used for qPCR were as follows:

*SAMMSON* Fw: CCTCTAGATGTGTAAGGGTAGT, Rv: TTGAGTTGCATAGTTGAGGAA

SDHA Fw: TGGGAACAAGAGGGCATCTG, Rv: CCACCACTGCATCAAATTCATG

HPRT1 Fw: TGACACTGGCAAAACAATGCA, Rv: GGTCCTTTTCACCAGCAAGCT

UBC Fw: ATTTGGGTCGCGGTTCTTG, Rv: TGCCTTGACATTCTCGATGGT

TBP Fw: CACGAACCACGGCACTGATT, Rv: TTTTCTTGCTGCCAGTCTGGAC

GAPDH Fw: TGCACCACCAACTGCTTAGC, Rv: GGCATGGACTGTGGTCATGAG

MALAT1 Fw: GGATCCTAGACCAGCATGCC, Rv: AAAGGTTACCATAAGTAAGTTCCAGAAAA

eGFP Fw: CAAGCAGAAGAACGGCATCA, Rv: AGGTAGTGGTTGTCGGGCA

### Differential gene expression analysis by RNA sequencing

RNA sequencing was performed on quadruplicates of NTC ASO (50 nM), *SAMMSON* ASO (50 nM), GDC-0349 (0.625 µM) and *SAMMSON* ASO (50 nM) + GDC-0349 (0.625 µM) treated 92.1 and OMM1 cells. Libraries for RNA sequencing were prepared using the Quant Seq 3’end library prep according to the manufacturer’s instructions (Lexogen, Vienna, Austria) using 2.5 µl input of RNA lysates and quantified on a Qubit Fluorometer prior to single-end sequencing with 75 bp read length on a NextSeq 500 sequencer (Illumina, San Diego, California, USA).

RNA sequencing data from UM patient derived xenograft (PDX) samples were obtained from Dewaele *et al*.(10) Libraries for RNA sequencing were prepared using the Truseq mRNA library prep according to the manufacturer’s instructions (Illumina) (500 ng input of purified RNA) and quantified on a Qubit Fluorometer prior to single-end sequencing with 75 bp read length on a NextSeq 500 sequencer (Illumina). Reads were mapped to the human genome (hg38) using STAR and gene expression was quantified using HTSeq (v0.6.1). Differentially expressed genes were identified using DESeq2 (v1.26.0).

A heatmap was created to visualize the differentially expressed genes in at least one condition irrespective of cell line differences. First, number of htseq raw counts >0 in at least half of the samples (per cell line) was used as a first selection, followed by differential expression analysis (design = ∼ cell line + treatment) using DESeq2 (v1.26.0). The DESeq2 results were normalized using normTransform (log2 (n+1)) and filtered per cell line. The input for the heatmap was scaled and clustered by rows with the centroid method using a Manhattan distance calculation.

### Statistical analysis

Statistical analyses and data visualizations were performed with Graphpad Prism version 9.2.0 (GraphPad Software, San Diego, California, USA). The individual data points and mean or mean ± s.d. are presented, unless otherwise specified. Significance of comparisons between more than two groups were determined using one-way ANOVA or two-way ANOVA with multiple testing correction. The level of statistical significance was set at p<0.05 (* p≤0.05, ** p≤0.01, *** p≤0.001, **** p≤0.0001).

## Results

### Compound screening reveals candidate synergistic combinations with SAMMSON inhibition

We performed an unbiased screen to identify compounds that, when combined with a lipid complexed *SAMMSON* inhibiting ASO, induce an additive or synergistic effect on uveal melanoma cell viability (Supplemental Fig 1). To that end, 2911 clinical stage compounds were combined with a lipid complexed *SAMMSON* inhibiting ASO or NTC ASO in the uveal melanoma cell line 92.1, followed by cell viability measurements and B-score data transformation (Supplemental Table 1). To compare the relative viability effect of each compound in the *SAMMSON* ASO and NTC ASO condition, the B-scores of the compounds in the *SAMMSON* ASO condition were subtracted from the B-scores in the NTC ASO condition. We selected 384 compounds for a confirmation screen based on negative B-scores in the *SAMMSON* ASO condition (lower relative viability in the *SAMMSON* ASO condition compared to all samples in the *SAMMSON* ASO condition) and positive B-score differences (lower relative viability in the *SAMMSON* ASO condition compared to the NTC ASO condition) (Fig 1 A and Supplemental Table 2). From the confirmation screen, 18 compounds were selected for further *in vitro* validation (Fig 1 B, C). Eleven of these compounds (i.e. clioquinol, fenofibric acid, LDN-193189, mifepristone, oltipraz, oxolinic acid, pimozide, propylthiouracil, ranitidine hydrochloride, riboflavin and INH-6) had no or only minor effects on cell viability as single agents, but further decreased cell viability when combined with *SAMMSON* ASO compared to *SAMMSON* ASO alone. The other 7 compounds (i.e. amsacrine, CI-994, ebastine, entinostat, fluphenazine, GDC-0349 and perphenazine) had a considerable effect on cell viability as single agents, but resulted in a substantial additional decrease on cell viability when combined with *SAMMSON* ASO treatment compared to the reductions obtained in the single compound or *SAMMSON* ASO treatment condition. These latter compounds were selected based on the median fold change differences between *SAMMSON* ASO and combined *SAMMSON* ASO - compound treated cells (data not shown). For each compound, whether or not combined with *SAMMSON* ASO, we mapped dose- and time-dependent effects in two UM cell lines (92.1 and OMM1) using cell viability measurements and time-lapse microscopy. We observed synergistic effects, calculated by means of excess over Bliss scores(34), for 6 compound - *SAMMSON* ASO combinations (i.e. amsacrine, GDC-0349, mifepristone, CI-994, entinostat and INH-6) (Fig 1 D, F and Supplemental Fig 2 A-C). The highest synergy scores for both UM cell lines could be obtained with mTOR inhibitor GDC-0349 (Fig 1 D-F), which we selected for further experiments. To confirm that the observed synergism resulted from mTOR inhibition, and not from GDC-0349 off-target effects, we evaluated additional mTOR inhibitors. Combining *SAMMSON* inhibition with temsirolimus, rapamycin and torkinib resulted in synergistic viability reductions comparable to GDC-0349 (Fig 2 A).

**Fig 1.**
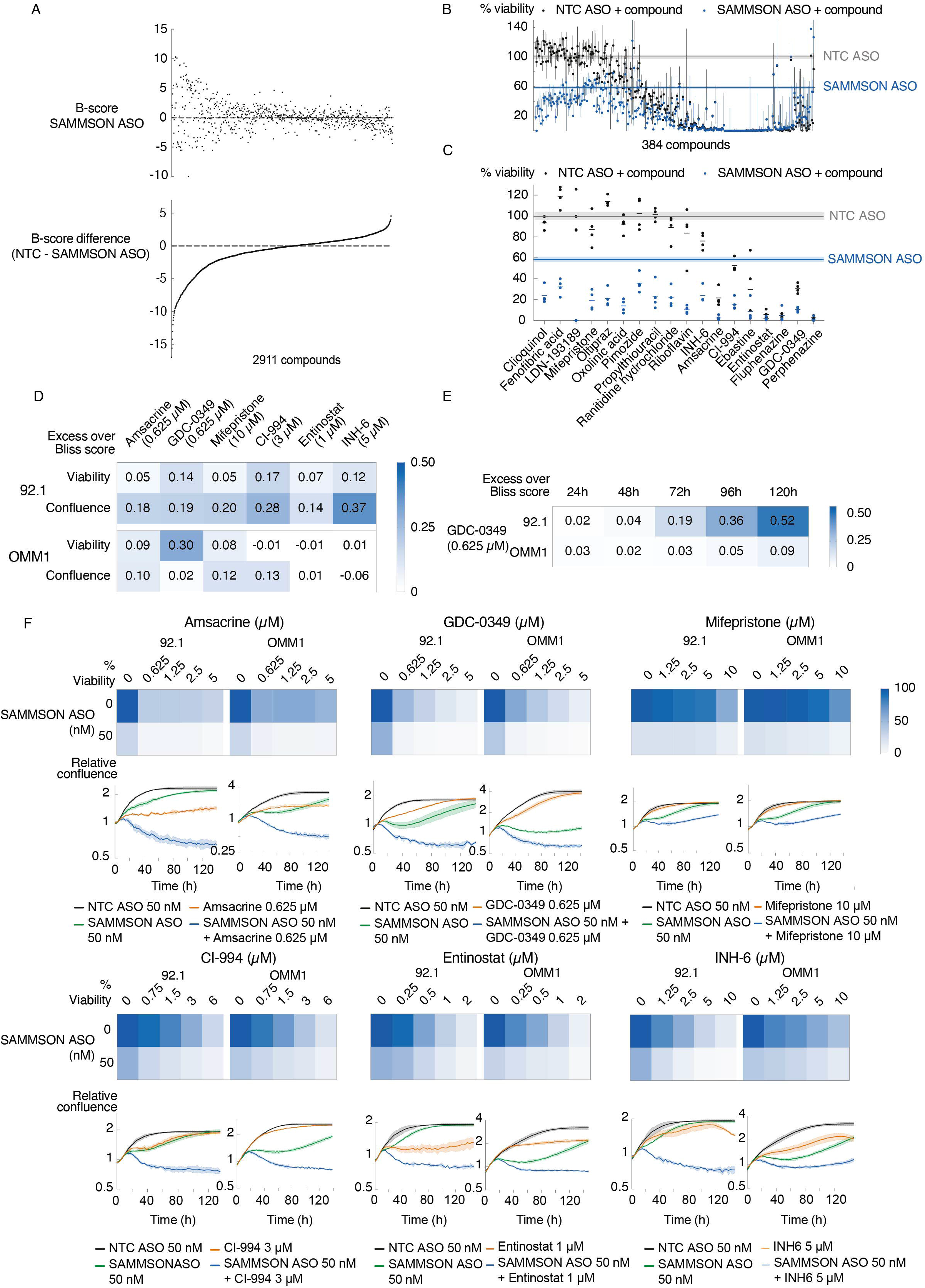
*In vitro* validation of *SAMMSON* inhibition in combination with clinical stage compounds. **A**. B-scores as a viability measure in UM cell line 92.1 48h after transfection with NTC ASO or *SAMMSON* ASO (35 nM) and compound treatment (10 µM). The upper panel presents the B-scores of the compounds in the *SAMMSON* ASO condition, the lower panel present the B-score difference (B-score NTC ASO - B-score *SAMMSON* ASO) of the same compound. **B, C**. Cell viability results 48h after treatment of UM cell line 92.1 treated with either NTC ASO or *SAMMSON* ASO (35 nM) whether or not in combination with a compound (10 µM). Values are relative to NTC ASO treated cells (grey horizontal line). The blue horizontal line represents the median viability effect of *SAMMSON* ASO treated cells. Black and blue dots are showing the median viability effects of each compound when combined with NTC ASO and *SAMMSON* ASO, respectively. Error bars represent ± 95% confidence interval (CI) of n=4. **C**. Detailed viability results of the 18 selected compounds from the confirmation screen (n=4). **D**. Excess over Bliss scores calculated 72h after treatment with compound and *SAMMSON* ASO (50 nM) and compared to both compound and *SAMMSON* ASO effect alone in 2 UM cell lines (92.1 and OMM1) with 2 different read outs (cell viability and confluence). In each treatment condition 3 replicates were included. **E**. Excess over Bliss scores calculated at different timepoints after treatment with GDC-0349 (0.625 µM) and *SAMMSON* ASO (50 nM) and compared to both GDC-0349 and *SAMMSON* ASO alone in 2 UM cell lines (92.1 and OMM1) with confluence as readout (n=3). **F**. Cell viability (72h time point, n=3) and relative confluence effects of UM cell lines 92.1 and OMM1 after treatment with NTC ASO or *SAMMSON* ASO (50 nM) with varying concentrations of 6 selected compounds. The real-time effect on confluence is shown for a fixed compound concentration. The confluence results are presented as the mean of 3 replicates ± s.d. and are relative to the first 4 measured data points.

**Fig 2.**
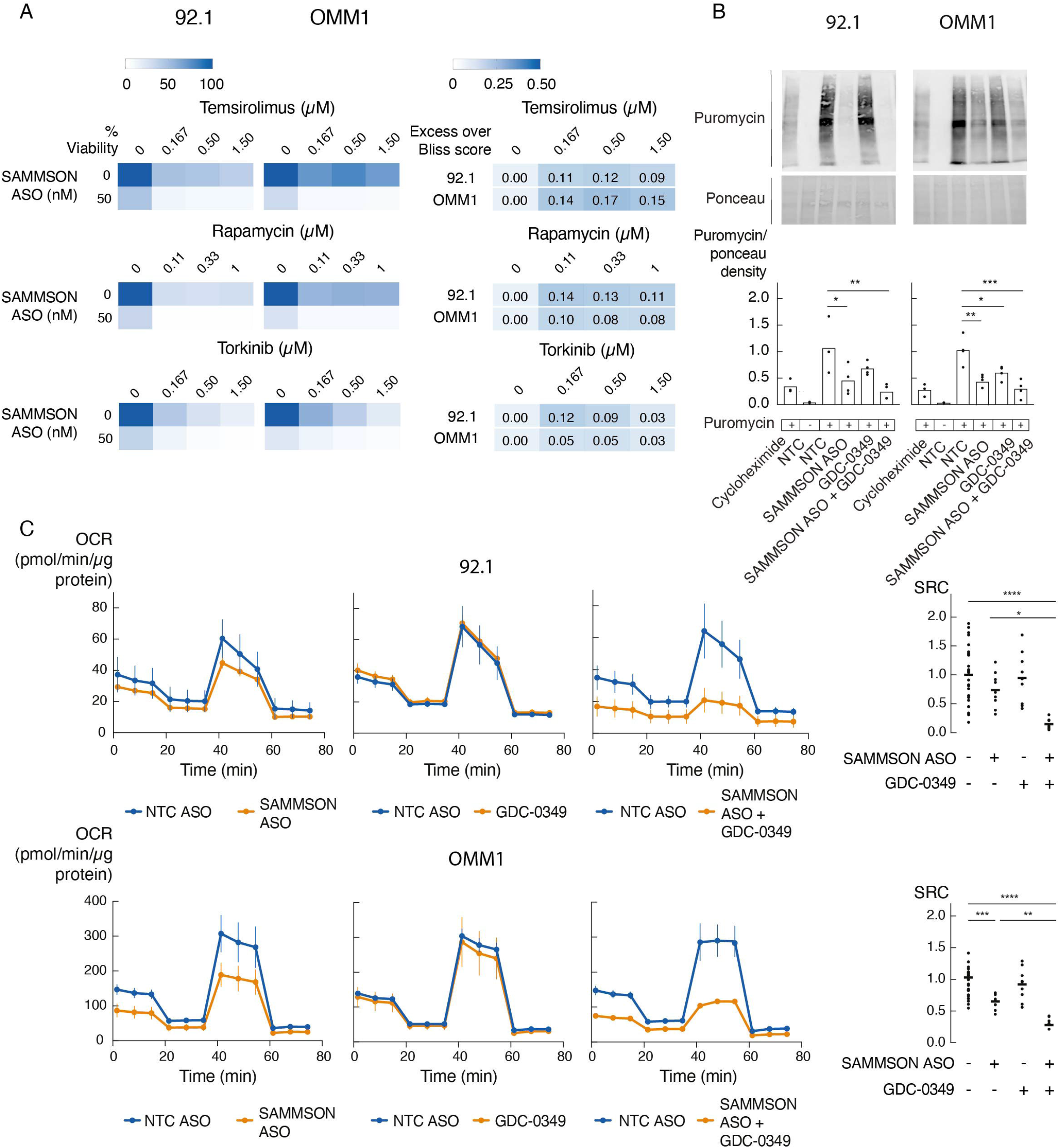
Combining mTOR inhibition with *SAMMSON* inhibition further reduces cell viability with additional effects on translation and mitochondrial function. **A**. Cell viability effects after 72h of treatment with varying concentration of mTOR inhibitors temsirolimus, rapamycin and torkinib in combination with *SAMMSON* ASO (50 nM) in UM cell lines 92.1 and OMM1 (n=3). The excess over Bliss scores were calculated for each compound concentration in both cell lines. **B**. Representative images of WB-SUnSET analysis of UM cells treated with cycloheximide (translation inhibitor, positive control), scrambled ASO (NTC, 50 nM) (without puromycin, negative control), NTC ASO (50 nM), *SAMMSON* ASO (50 nM), GDC-0349 (0.625 µM) or the combination of *SAMMSON* ASO (50 nM) and GDC-0349 (0.625 µM). Quantification of protein synthesis measured by calculating the intensity of the puromycin signal on WB. The individual data points (n=3) and mean are presented. P-values were calculated using two-way ANOVA with Tukey’s multiple comparisons test. **C**. Oxygen Consumption Rate (OCR) measurements over time after sequential injections of oligomycin, fluoro-carbonyl cyanide phenylhydrazone (FCCP) and rotenone/antimycin A in UM cell lines treated for 24h with NTC ASO (50 nM), *SAMMSON* ASO (50 nM), GDC-0349 (0.625 µM) or the combination of *SAMMSON* ASO (50 nM) and GDC-0349 (0.625 µM). Data are represented as the mean of 3 replicates ± s.d. Spare respiratory capacity (SRC) was obtained by subtracting the basal respiration from the maximal respiration. The individual data points and mean are presented. P-values were calculated using one-way ANOVA with Tukey’s multiple testing correction. * p≤0.05, ** p≤0.01, *** p≤0.001, **** p≤0.0001.

### mTOR inhibitor GDC-0349 enhances the molecular and cellular phenotype induced by SAMMSON knockdown

As we have described previously(10), *SAMMSON* is essential for the survival of both primary and metastatic uveal melanoma tumor cells through its involvement in cytosolic and mitochondrial translation. *SAMMSON* inhibition consequently impairs mitochondrial function, which phenotypically results in a reduction of cell viability and induction of apoptosis. We therefore, evaluated the impact on protein synthesis and mitochondrial function when combining *SAMMSON* and mTOR inhibition. Quantification of global translation levels after GDC-0349 treatment using a puromycin incorporation assay (SUnSET(35)) showed impaired translation rates of 42% in OMM1 and 38% in 92.1 (Fig 2 B and Supplemental Fig 3, p=0.0187 (OMM1) and n.s. (92.1), one-way ANOVA). In line with previous results(10), *SAMMSON* ASO reduced the translation rates with 56% and 61%, respectively (Fig 2 B and Supplemental Fig 3, p=0.0017 (OMM1) and p=0.0259 (92.1), one-way ANOVA). When combining GDC-0349 and *SAMMSON* ASO treatments, an additional reduction in translational rate of 36% in OMM1 and 47% in 92.1 compared to *SAMMSON* ASO alone was observed which finally results in a total reduction of 73% and 82% compared to NTC ASO treatment, respectively (p=0.0001 (OMM1) and p=0.0023 (92.1), one-way ANOVA). To further investigate the effect on mitochondrial function, we measured the differences in oxygen consumption rate (OCR) after injections of oligomycin, FCCP and rotenone/antimycin A(36). Treatment of UM cells with *SAMMSON* ASO and mTOR inhibitor GDC-0349 24h prior to OCR measurement resulted in a significant decrease in mitochondrial spare respiratory capacity (SRC) of 72% and 85% compared to NTC ASO treatment in OMM1 and 92.1, respectively (Fig 2 C, p<0.0001, one-way ANOVA). Compared to *SAMMSON* ASO treatment, the combined treatment resulted in an additional decrease in OCR of 57% and 80% (p=0.0052 (OMM1) and p=0.0203 (92.1), one-way ANOVA).

### mTOR inhibitor GDC-0349 enhances intracellular ASO uptake and activity

To elucidate the mechanism underlying the synergistic effect of combined *SAMMSON* and mTOR inhibition, we performed shallow RNA sequencing of UM cell lines 92.1 and OMM1 treated with either NTC ASO, *SAMMSON* ASO, GDC-0349 or the combination of *SAMMSON* ASO and GDC-0349. Differential gene expression analysis revealed very few differentially expressed genes (DEG) in the GDC-0349 treated UM cell lines (28 DEGs (12 up- and 16 downregulated) in 92.1 and 6 DEGs (4 up- and 2 downregulated) in OMM1) (adjusted p-value <0.05). In comparison, *SAMMSON* ASO treatment resulted in 312 DEGs (202 up- and 110 downregulated) in 92.1 and 301 DEGs (208 up- and 93 downregulated) in OMM1. When combining both *SAMMSON* ASO and GDC-0349, the amount of DEGs significantly increased to 1723 DEGs (769 up- and 954 downregulated) in 92.1 and 887 DEGs (480 up- and 407 downregulated) in OMM1. Notably, a substantial fraction of DEGs in the *SAMMSON* ASO treated condition (68% (213/312) in 92.1 and 86% (258/301) in OMM1) overlapped with the DEGs in the combined treatment, with more pronounced fold changes in the combined treatment (Fig 3 A). A heatmap visually representing 341 genes that are differentially expressed (adjusted p-value ≤0.001 and absolute(log2FoldChange) > 1) in at least one treatment condition in both cell lines illustrates that GDC-0349 further enhances the differential gene expression profile observed upon *SAMMSON* inhibition (Fig 3 B). These observations suggested that mTOR inhibition further reduced *SAMMSON* expression in ASO treated cells. While the *SAMMSON* ASO reduced *SAMMSON* expression with 63% and 28% in OMM1 and 92.1, respectively (Fig 4 A, p≤0.0001 (OMM1) and p=0.0001 (92.1), one-way ANOVA), the combination with GDC-0349 further reduced *SAMMSON* expression up to 78% and 49% (p<0.0001, one-way ANOVA). We again evaluated if alternative mTOR inhibitors had similar effects. The addition of mTOR inhibitor temsirolimus or rapamycin to *SAMMSON* ASO resulted in an additional decrease in *SAMMSON* expression of 82% and 73% in OMM1 and 39% and 48% in 92.1, respectively, compared to *SAMMSON* ASO monotherapy (Fig 4 B, p<0.0001 (OMM1), p=0.0009 (92.1, temsirolimus) and p=0.0343 (92.1, rapamycin), one-way ANOVA).

**Fig 3.**
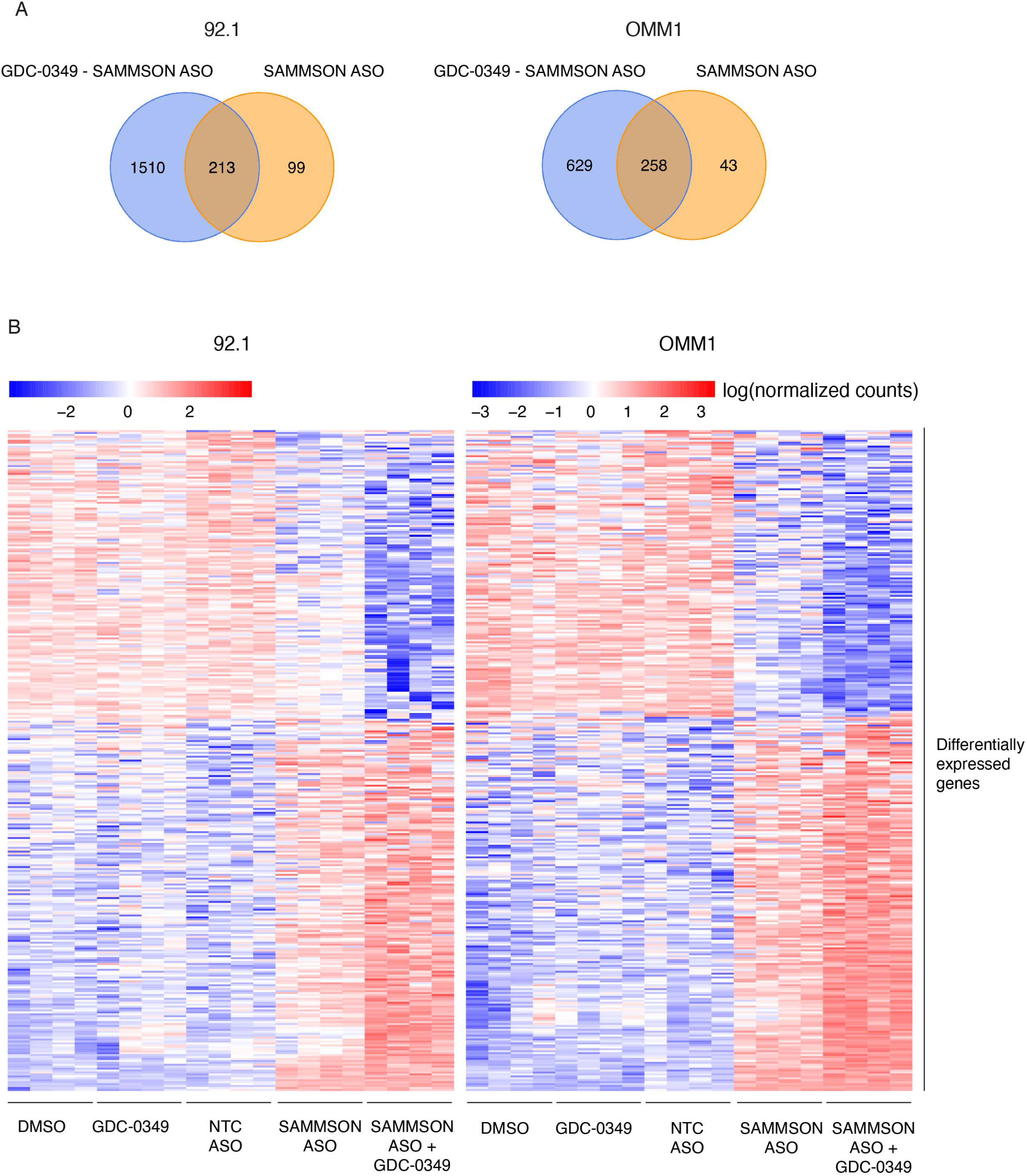
mTOR inhibitor GDC-0349 enhances differential gene expression profile of *SAMMSON* inhibition. **A**. Venn diagram presenting the total number and overlapping number of DEGs in the *SAMMSON* ASO (50 nM) and combined *SAMMSON* ASO (50 nM) and GDC-0349 (0.625 µM) condition compared to NTC ASO in UM cell lines 92.1 and OMM1. **B**. Heatmap for each cell line presenting the genes that are differentially regulated in at least one condition (n=341).

**Fig 4.**
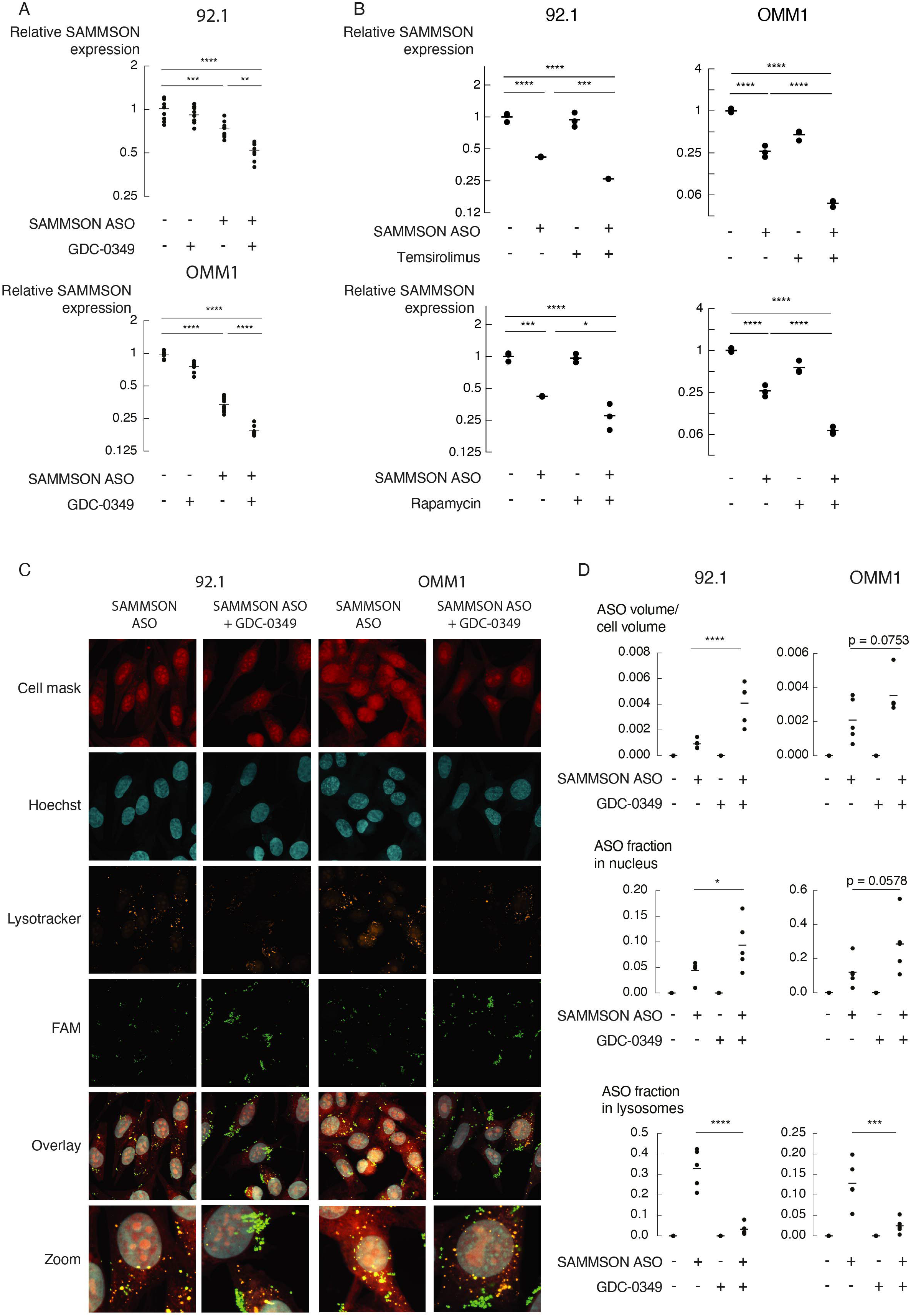
mTOR inhibition enhances intracellular ASO uptake, activity and consequently target knockdown, **A, B.** Relative *SAMMSON* expression 48h after transfection in UM cell lines 92.1 and OMM1 transfected with 50 nM of a scrambled ASO (NTC) or *SAMMSON* ASO whether or not combined with mTOR inhibitors GDC-0349 (A) (0.625 µM, n=9), temsirolimus (B) (0.5 µM, n=3) or rapamycin (B) (0.33 µM, n=3). The individual data points and mean are presented. **C**. Confocal microscopy images of UM cell lines 92.1 and OMM1 transfected with *SAMMSON* ASO (50 nM) and the combination of *SAMMSON* ASO (50 nM) and GDC-0349 (0.625 µM) for 24h. Representative images of the cells (cell mask), nucleus (Hoechst), lysosomes (lysotracker) and FAM-labeled *SAMMSON* ASO (FAM). Yellow spots in the overlay and zoom images represent ASO localization to lysosomes. **D**. Relative ASO volume inside the cell to the total cell volume and the fraction in the nucleus and lysosomes when treated with either *SAMMSON* ASO, GDC-0349 or the combination of both. The individual data points of 5 randomly selected fields per condition (containing ≥ 30 cells in total per condition) and mean are presented. P-values in A, B and D were calculated using one-way ANOVA with Tukey’s multiple testing correction. * p≤0.05, ** p≤0.01, *** p≤0.001, **** p≤0.0001.

Based on these observations, and a recently published report of Ochaba J. et al.(37), we hypothesized that mTOR inhibition promotes intracellular uptake and trafficking of the *SAMMSON* ASO-lipid complexes in UM cells, resulting in a higher intracellular ASO concentration and enhanced *SAMMSON* knockdown. To test this hypothesis, we transfected UM cells with a 3’ FAM labeled *SAMMSON* ASO, whether or not in combination with GDC-0349. Cells were stained with a lysosome selective dye (LysoTracker Red DND-99) to examine the quantity and intracellular localization of the fluorescently labeled ASO. Confocal imaging revealed that GDC-0349 treatment induces a 1.7- and 4.5-fold increase in ASO quantity per cell (Fig 4 C, D, p=0.0753 (n.s., OMM1) and p<0.0001 (92.1), one-way ANOVA) and a 2.4- and 2.1-fold increase in ASO quantity in the nucleus (Fig 4 C, D, p=0.0578 (n.s., OMM1) and p=0.0409 (92.1), one-way ANOVA) compared to cells that were not supplemented with GDC-0349. Furthermore, GDC-0349 significantly decreased the amount of *SAMMSON* ASO in the lysosomes by as much as 80% and 90% in OMM1 and 92.1, respectively (Fig 4 C, D, p=0.0002 (OMM1) and p<0.0001 (92.1), one-way ANOVA).

To further validate these observations, we treated UM cells with a MALAT1 ASO and GAPDH ASO, whether or not in combination with GDC-0349. Transfection of MALAT1 ASO and GAPDH ASO significantly decreased *MALAT1* and *GAPDH* expression with 92% and 86% in OMM1 and 45% and 42% in 92.1, respectively (Fig 5 A, p<0.0001 (OMM1), p=0.0002 and p=0.0003 (92.1), one-way ANOVA). Addition of GDC-0349 further reduced *MALAT* expression levels with 68% in OMM1 and *GAPDH* expression levels with 64% and 49% in both OMM1 and 92.1, respectively (Fig 5 A, p<0.0001, one-way ANOVA). Taken together, our results demonstrate that the mTOR inhibitor GDC-0349 enhances uptake and reduces lysosomal accumulation of ASO-lipid complexes in UM cells, significantly improving target knockdown efficiency.

**Fig 5.**
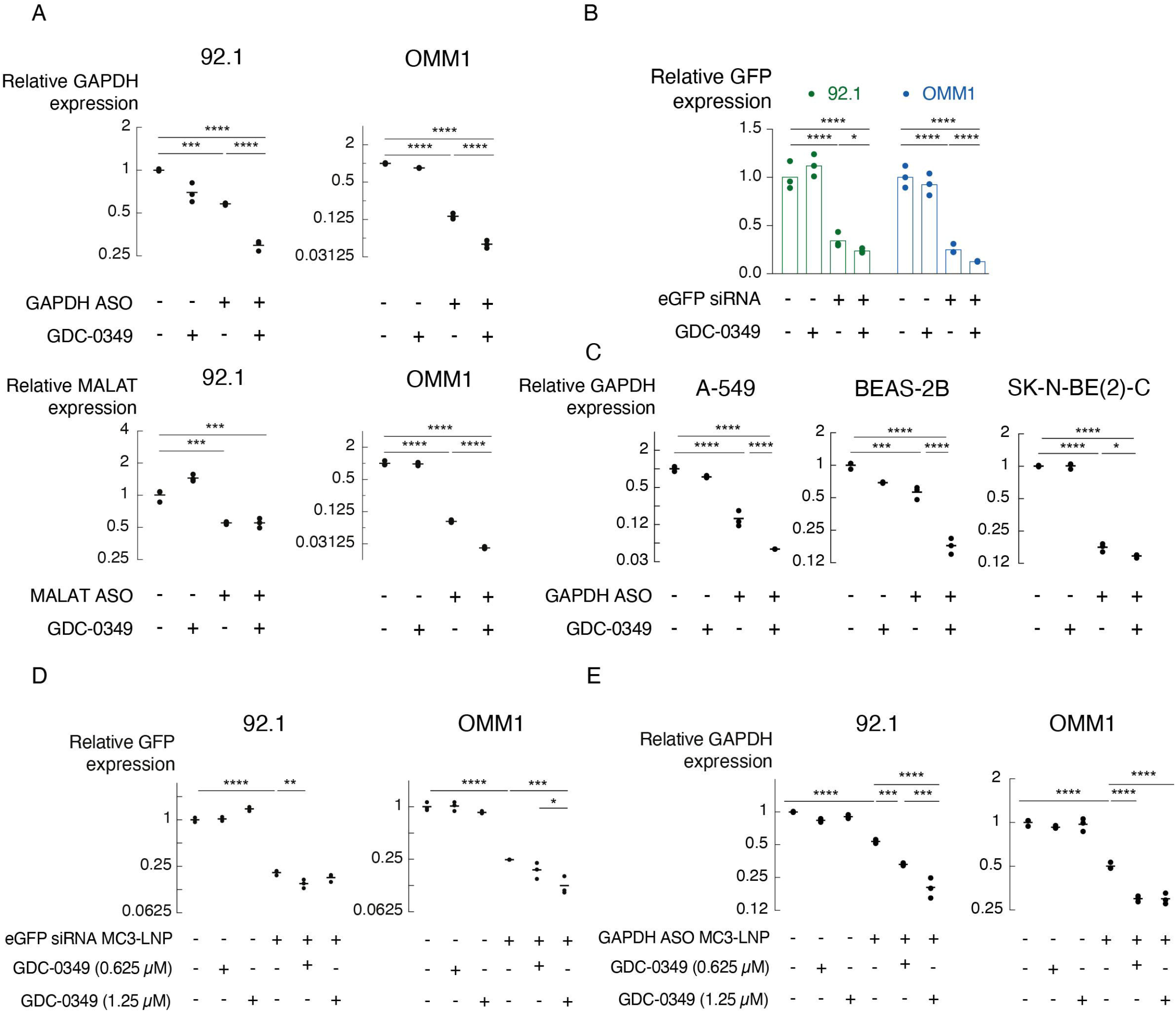
Improved ASO and siRNA target knockdown when combined with mTOR inhibition in multiple cell types and independent of oligonucleotide formulation. **A**. Relative *GAPDH* and *MALAT* expression 48h after transfection in UM cell lines 92.1 and OMM1 transfected with 50 nM of a scrambled ASO, GAPDH or MALAT ASO (50 nM) and whether or not combined with GDC-0349 (0.625 µM). The three individual data points and mean are presented. **B**. Relative *GFP* expression in 92.1 and OMM1 upon transfection for 48h with scrambled siRNA or eGFP siRNA (100 nM) whether or not in combination with mTOR inhibitor GDC-0349 (0.625 µM). The three individual data points and mean are presented. **C**. Relative *GAPDH* expression in A-549 lung adenocarcinoma cells, BEAS-2B bronchial epithelial cells and SK-N-BE(2)-C neuroblastoma cells upon treatment for 48h with NTC ASO or GAPDH ASO (50 nM) and GDC-0349 (0.625 µM) or DMSO. The three individual data points and mean are presented. **D-E**. Relative *GFP* (D) and *GAPDH* (E) expression upon treatment for 48h with NTC siRNAs or ASOs, eGFP siRNAs (D) or GAPDH ASOs (E) in MC3-LNP formulation. The effects upon combination with two concentrations of GDC-0349 (0.625 µM and 1.25 µM) and DMSO are shown for UM cell lines 92.1 and OMM1. The three individual data points and mean are presented. P-values in A, C, D and E were calculated using one-way ANOVA with Tukey’s multiple testing correction. P-values in B were calculated using two-way ANOVA with Tukey’s multiple testing correction. * p≤0.05, ** p≤0.01, *** p≤0.001, **** p≤0.0001.

### mTOR inhibition improves delivery of oligonucleotides independent of formulation and cell type

We then evaluated if the improved oligonucleotide delivery in UM cells upon mTOR inhibition also applies to double stranded siRNAs. To that purpose, we treated OMM1 and 92.1 cells stably expressing GFP with a GFP siRNA. Transfection of GFP siRNA significantly decreased GFP expression with 75% and 66% in both OMM1-GFP and 92.1-GFP, respectively (Fig 5 B, p<0.0001, two-way ANOVA). Combining GFP siRNA with GDC-0349 further reduced GFP expression with 49% and 69% (Fig 5 B, p<0.0001 (OMM1) and p=0.0238 (92.1), two-way ANOVA), suggesting that mTOR inhibition improves delivery of both ASOs and siRNAs to UM cells.

To investigate if mTOR inhibition also improves ASO delivery in other cell types, we transfected A-549 lung adenocarcinoma cells, BEAS-2B bronchial epithelial cells and SK-N-BE(2)-C neuroblastoma cells with a GAPDH ASO whether or not in combination with GDC-0349. A significant decrease in *GAPDH* expression was obtained in all three cell lines (84% in A-549 (p<0.0001), 44% in BEAS-2B (p=0.0005) and 82% in SK-N-BE(2)-C (p<0.0001)), which was further enhanced with 68% in A-549 and BEAS-2B and 17% in SK-N-BE(2)-C when combining with GDC-0349 (Fig 5 C, p<0.0001 (A-549 and BEAS-2B), p=0.0144 (SK-N-BE(2)-C), one-way ANOVA).

Finally, we investigated if the impact of mTOR inhibition can be extended to alternative oligonucleotide formulations using a more advanced lipid nanoparticle (LNP) formulation approved for clinical application. More specifically, we generated LNP-encapsulated GFP siRNAs (eGFP siRNA MC3-LNP) and GAPDH ASOs (GAPDH ASO MC3-LNP) using (6Z,9Z,28Z,31Z)-heptatriaconta-6,9,28,31-tetraen-19-yl4-(dimethylamino)butanoate (DLin-MC3-DMA) ionizable amino-lipids(21, 26, 38). Administration of 2.5 nM of eGFP siRNA MC3-LNP significantly reduced *GFP* expression levels with 76% and 79% in both OMM1-GFP and 92.1-GFP, respectively (Fig 5 D, p<0.0001, one-way ANOVA). When combined with 0.625 µM of GDC-0349, *GFP* expression levels were further reduced by 24% and 28% (Fig 5 D, n.s. (OMM1-GFP) and p=0.0059 (92.1-GFP). Increasing the GDC-0349 concentration to 1.25 µM resulted in a *GFP* expression reduction of 34% compared to eGFP siRNA MC3-LNP in OMM1-GFP cells (Fig 5 D, p=0.0362 (OMM1-GFP), one-way ANOVA, no additional reduction observed in 92.1-GFP). Comparable results were obtained with GAPDH ASO MC3-LNP where a reduction of 50% and 46% in *GAPDH* expression in OMM1 and 92.1 was obtained with 20 nM of GAPDH ASO MC3-LNP (Fig 5 E, p<0.0001, one-way ANOVA). Addition of 0.625 µM of GDC-0349 further reduced the *GAPDH* expression with 40% and 39%, respectively (Fig 5 E, p<0.0001 (OMM1) and p=0003 (92.1), one-way ANOVA). When increasing the GDC-0349 concentration to 1.25 µM, an additional reduction in *GAPDH* expression of 38% compared to 0.625 µM of GDC-0349 was obtained in 92.1 cells (Fig 5 E, p=0.0002 (92.1), one-way ANOVA, no additional reduction observed in OMM1).

In summary, these results demonstrate that mTOR inhibition enhances delivery of ASOs and siRNAs in UM and other cell types, using both cationic delivery (lipofectamine) and MC3-LNPs (DLin-MC3-DMA).

## Discussion

Despite the effective treatment modalities of the primary tumor, uveal melanoma is characterized by a high rate of liver metastasis for which no effective treatments are available, resulting in a median survival time of less than 12 months(7). Any treatment option that could control dissemination, would potentially increase the survival time of UM patients. We recently identified the lncRNA *SAMMSON* as a lineage survival gene expressed in >80% of uveal melanoma tumors with higher expression levels in metastatic UM tumors compared to primary UM tumors(10). ASO-mediated silencing of *SAMMSON* induces a potent anti-tumor response both *in vitro* and *in vivo*, demonstrating a potential treatment strategy to treat both primary and metastatic UM patients. In order to enhance the therapeutic potential, we screened 2911 clinical stage compounds in combination with *SAMMSON* ASO to identify synergistic cell viability interactions. It’s worth noting that, at the dose used for compound screening (10 µM), several compounds were too toxic to be able to observe additive or synergistic effects, which is a limitation of the current study. It would be beneficial to screen multiple ASO and compound concentrations in the initial screen to reveal all interesting combinations. A selection of 18 compounds was extensively validated in two UM cell lines where multiple concentrations were tested at different time points using various read outs. For some compounds, we failed to reproduce the screening results, which might be due to differences in chemical and physical quality between compounds in the screening library and compounds used for validation and differences in the automated liquid handling and signal detection methods between screening and validation experiments.

Our screening results revealed mTOR inhibitor GDC-0349 as a synergistic compound with *SAMMSON* ASO in UM. mTOR activity is frequently deregulated in various diseases including Alzheimer’s disease, diabetes and human cancers such as breast cancer, lung carcinomas and uveal melanoma(39, 40). Hence, mTOR is a promising therapeutic target for cancer treatment. mTOR is the catalytic subunit of two distinct complexes, mTOR complex 1 (mTORC1) and 2 (mTORC2), which differ in their composition, downstream targets and pathways. mTORC1 is involved in protein synthesis and anabolic processes including lipid and nucleotide synthesis, while mTORC2 is involved in cytoskeleton organization, cell survival and regulation of glucose and lipid metabolism(41). GDC-0349 inhibits both mTOR complexes(42, 43), but alternative mTOR inhibitors targeting only mTORC1 (i.e. rapamycin and temsirolimus) also resulted in synergistic effects when combined with *SAMMSON* ASO, suggesting mTORC1 to be the main complex involved. In our previous study, we demonstrated impaired cytosolic and mitochondrial translation, which consequently affects mitochondrial function when *SAMMSON* is inhibited in UM(10). When combined with mTOR inhibitor GDC-0349, the global translation rates and mitochondrial function are further reduced, although GDC-0349 monotherapy only affects translation without influencing mitochondrial function. Transcriptome analysis also revealed very few changes in gene expression after GDC-0349 treatment, suggesting that, at the applied concentration, GDC-0349 has a limited effect in UM cells. However, when combining mTOR inhibition with *SAMMSON* inhibition, the amount and the log fold change of DEGs was enhanced compared to SAMMON inhibition alone. In line with these observations, *SAMMSON* expression levels were further reduced in the combined treatment compared to *SAMMSON* ASO alone, which could also be confirmed with additional mTOR inhibitors. Enhanced target knockdown efficiencies were also observed with additional ASOs, such as MALAT and GAPDH ASO, suggesting this to be a general phenomenon. A recent study has shown enhanced knockdown efficiencies when combining mTOR inhibition with unformulated ASOs in free-uptake compatible cells(37). The increased ASO activity was associated with enhanced localization of ASOs to the autophagosomes, which contributes to ASO trafficking and facilitates lysosomal escape. Since the UM cells in this study are free-uptake incompatible cells, ASO delivery requires a lipid based formulation. The positively charged cationic liposomes interact electrostatically with the negatively charged nucleic acid sequences, resulting in the formation of a positively charged complex which is attracted to the negatively charged cell membrane and enters the cell through endocytosis(44, 45). It has been reported that cationic lipid formulations, such as lipofectamine 2000, induce autophagy in mammalian cells, hereby avoiding metabolic degradation into lysosomal compartments(46–48). In this study, we show that the amount of intracellular *SAMMSON* ASO, when delivered using a cationic lipid, is enhanced when combined with mTOR inhibition. Moreover, we observed a shift of ASO molecules in the lysosomal compartment towards the nucleus, explaining the enhanced ASO activity and target knockdown. Additional studies are required to further elucidate if mTOR inhibition in combination with cationic lipid ASO formulations reduce lysosomal accumulation through autophagy induction. To demonstrate the broad relevance of mTOR inhibition in nucleic acid delivery and activity *in vitro*, enhanced target knockdown efficiencies were obtained for siRNAs in UM cells and ASOs in various cancer cell types (lung adenocarcinoma and neuroblastoma) and a normal cell type (bronchial epithelial cells). In UM, GDC-0349 also improved target knockdown for more advanced lipid nanoparticle formulated ASOs and siRNAs containing ionizable amino lipids (MC3-LNP), which are the leading system for intravenously administered RNAi based therapeutics. MC3-LNP formulated ASOs and siRNAs are protected from degradation by endogenous nucleases, have a prolonged circulation time, reduced immune stimulation and accumulate mainly in the liver(49, 50). The latter might be beneficial for the treatment of metastatic UM tumors which preferably disseminate to the liver. Additional *in vivo* studies are required to further assess the potential of mTOR inhibition for improved target knockdown in UM tumor cells.

A combination of drugs can be called synergistic when the cellular effect induced in the combined treatment is larger than the summed responses of the individual drugs(51, 52). Instinctively, we expected the combination identified in this study to be catalogued as additive, since the identified underlying mechanism of the GDC-0349 – *SAMMSON* ASO combination does not support the concept of synergism. As 0.625 µM of GDC-0349 only has a negligible effect on cell viability, while the combination of *SAMMSON* ASO and GDC-0349 increases the cell viability reduction compared to *SAMMSON* ASO alone, this interaction could, however, be catalogued as synergistic by the Bliss Independence method. This demonstrates the importance of mechanistic insight underlying combination treatments to assess the output of synergistic calculation methods.

In conclusion, our work shows that mTOR inhibition improves lipid mediated uptake and reduces lysosomal accumulation of both ASOs and siRNAs in UM cells and other cell types. Consequently, mTOR inhibition enhances ASO- or siRNA-mediated target knockdown.

## Data availability

RNA-seq data will be deposited to Gene Expression Omnibus (GEO) database.

## Funding

This work was supported by a Foundation Against Cancer grant (Stichting tegen Kanker, FAF-C/2016/749), a Standup Against Cancer grant (Kom op tegen Kanker, 365C8916), a Ghent University Industrial Research Fund (IOF, F2015/IOF-Advanced/040), a Ghent University GOA research grant and Fund for Scientific Research Flanders (FWO Vlaanderen, G029317N). S.D., B.B. and L.D. are recipients of a grant from the Fund of Scientific Research Flanders (FWO Vlaanderen, 1149819N).

## Competing interests

No potential conflicts of interest were disclosed.

## Acknowledgments

The authors would like to thank Prof. Aart Jochemsen for providing the UM cell lines, the VIB Screening Core (Ghent, Belgium) for performing the compound screen and the VIB Bioimaging Core (Ghent, Belgium) for performing confocal imaging.

## Authors’ Contributions

P.M. conceived and supervised the project; S.D, L.D., B.D.P., B.B., N.Y, J.N. and S.E. designed and performed experiments; P.M., S.D., J.A analyzed the data; L.D. and S.E. made and provided the GFP constructs, R.M. produced virus particles for the modification of cell lines, B.B. and K.R. made, characterized and provided the siRNA and ASO MC3-LNP particles, K.R., R.V.C., S.E. contributed technical support and resources; S.D. and P.M wrote the paper. All authors contributed to manuscript editing and approved the final draft.

## Figure legends

**Supplemental Fig 1. Overview screening procedure**. *SAMMSON* ASO or NTC ASO transfection was combined with 2911 clinical stage compounds for an initial screen in UM cell line 92.1, followed by a selection of 384 compounds for a confirmation screen. Based on cell viability results, 18 promising compound-*SAMMSON* ASO combinations were validated in UM cell lines 92.1 and OMM1 using multiple doses, timepoints and read-out methods to finally select one compound for further mechanistic studies.

**Supplemental Fig 2. *In vitro* validation of *SAMMSON* inhibition in combination with the additional 12 clinical stage compounds. A-B**. Cell viability (72h time point, n=3) (A) and relative confluence (B) effects of UM cell lines 92.1 and OMM1 after treatment with NTC ASO or *SAMMSON* ASO (100 nM) with a fixed concentration of the additional 12 compounds. The confluence results are presented as the mean of 3 replicates ± s.d. and are relative to the first 4 measured data points. **C**. Excess over Bliss scores calculated 72h after treatment with compound and *SAMMSON* ASO (100 nM) and compared to both compound and *SAMMSON* ASO effect alone in UM cell lines 92.1 and OMM1 with 2 different read outs (cell viability and confluence). In each treatment condition 3 replicates were included.

**Supplemental Fig 3. Uncropped images of WB-SUnSET analysis**. Uncropped images of WB-SUnSET analysis of UM cells 92.1 and OMM1 treated with cycloheximide (translation inhibitor, positive control), scrambled ASO (NTC) (without puromycin, negative control), NTC ASO (50 nM), *SAMMSON* ASO (50 nM), GDC-0349 (0.625 µM) or the combination of *SAMMSON* ASO (50 nM) and GDC-0349 (0.625 µM).

**Supplemental Table 1. B-scores of 2911 ASO-compound combinations in the initial screen**.

**Supplemental Table 2. Cell viability results of 384 ASO-compound combinations in the confirmation screen**.

## Notes

### Competing Interest Statement

The authors have declared no competing interest.

