## Supplementary figures and images for "mTOR inhibition enhances delivery and activity of antisense oligonucleotides in uveal melanoma cells"

### Supplemental Fig 1

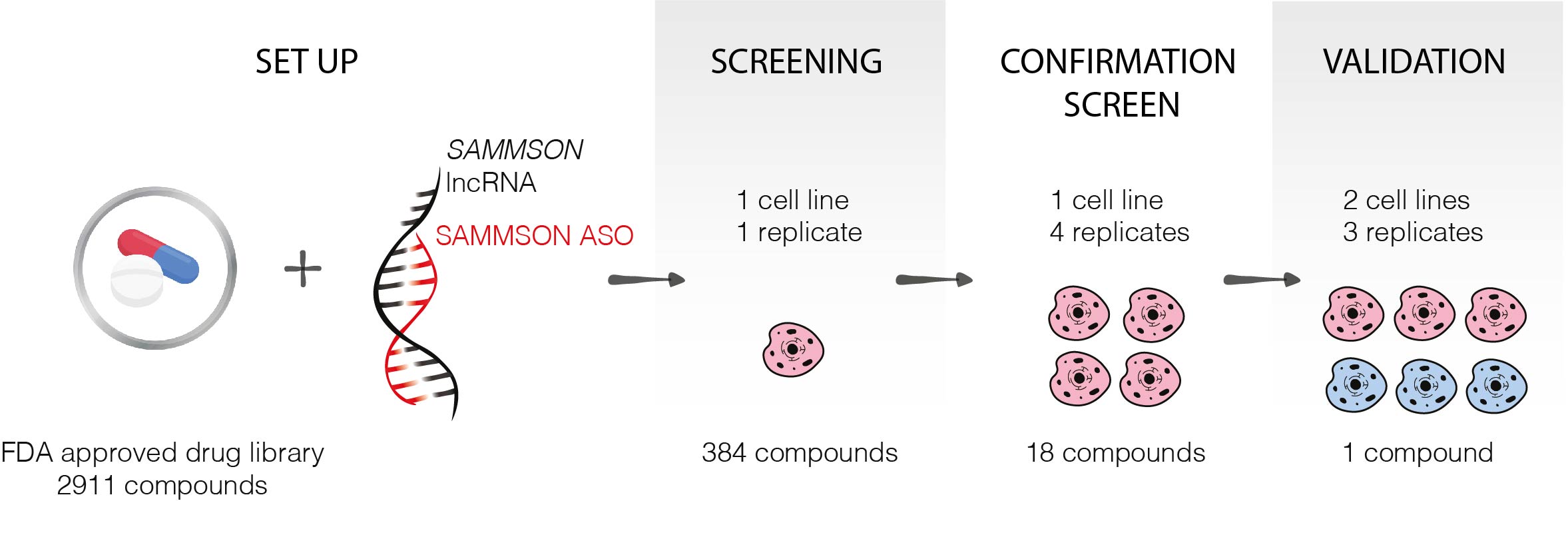

### Supplemental Fig 2

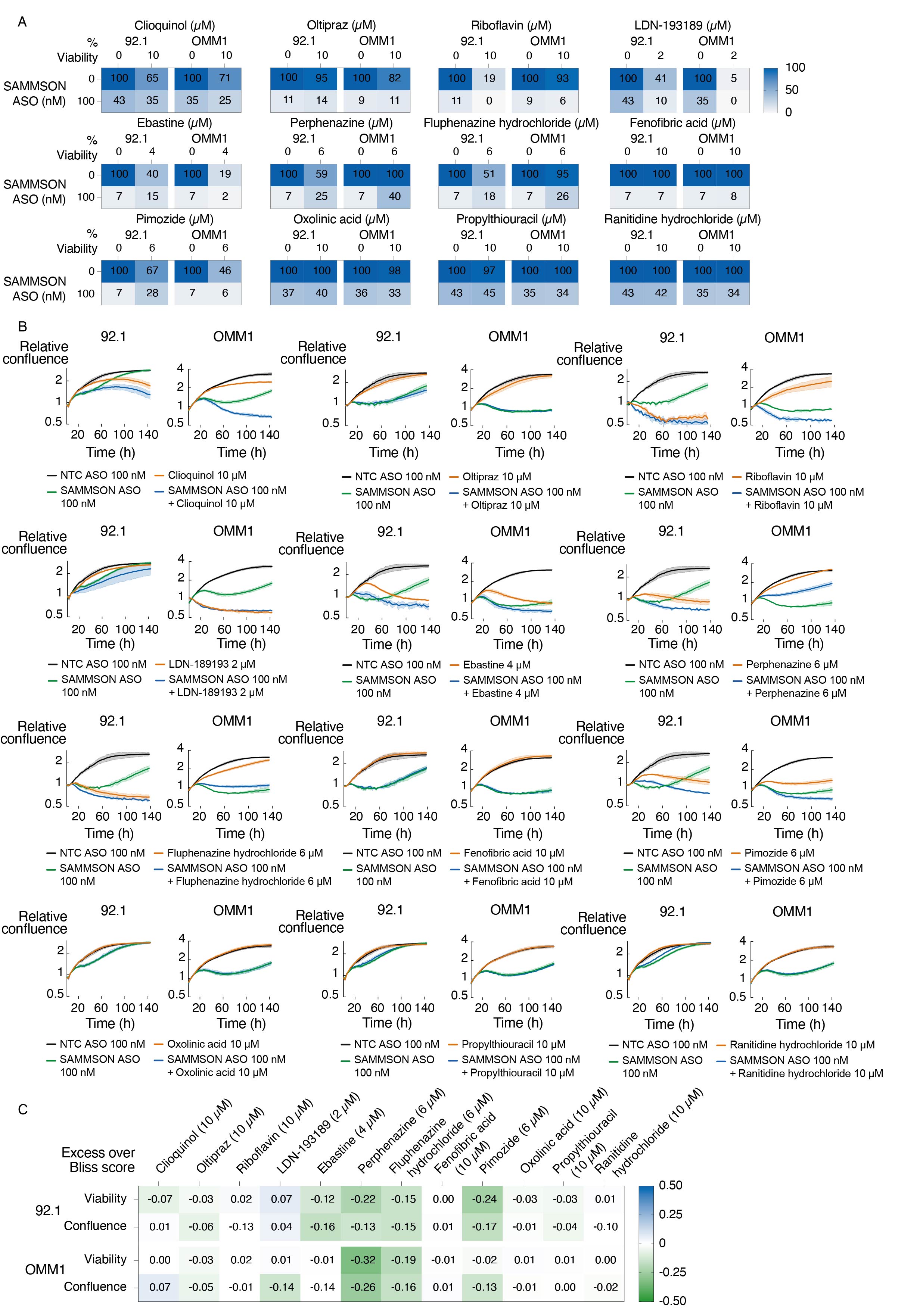

### Supplemental Fig 3

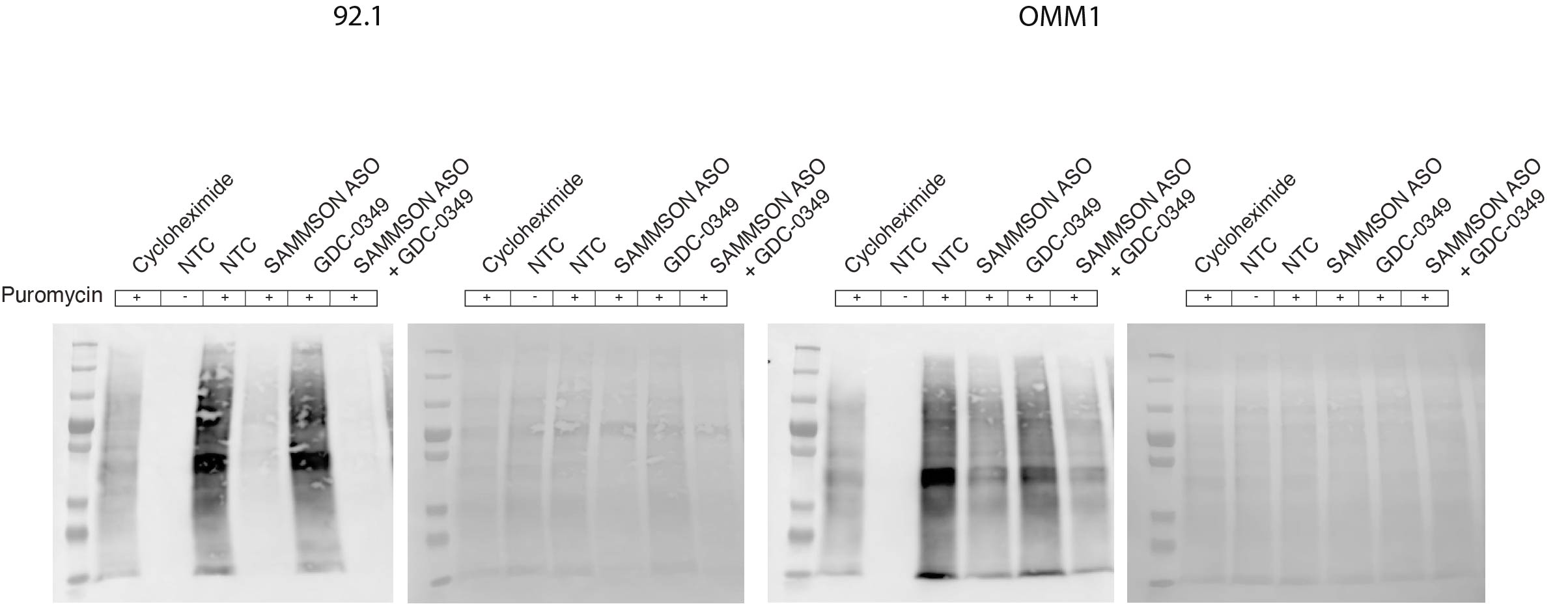
